# Ontogenetic variation in composition and bioactivity of common adder (*Vipera berus*) venom revealed by genome-guided proteomics and in vitro functional assays

**DOI:** 10.64898/2026.02.09.704791

**Authors:** Lennart Schulte, Johanna Eichberg, Alfredo Cabrera-Orefice, Harry Wölfel, Paul Hien, Maik Damm, Tilman Schell, Charlotte Gerheim, Alexander Ben Hamadou, Sabine Hurka, Thomas Lindner, Gregor Geisler, Ignazio Avella, Kornelia Hardes, Carola Greve, Tim Lüddecke

## Abstract

Ontogenetic shifts in diet are well documented in snakes and are increasingly linked to age-related venom variation. The common adder, *Vipera berus*, exhibits a dietary transition from predominantly ectothermic prey in its early life to increasingly incorporating endothermic prey as an adult. Here, we investigate whether this dietary shift is reflected in age-related changes in the venom composition and bioactivity of *V. berus*. Venoms from captive-bred *V. berus* from Germany were obtained and pooled across five age groups, from neonates to adults. Venom profiles were assessed by SDS-PAGE and genome-guided shotgun proteomics, with quantification based on normalized spectral abundance factors (NSAF) using a toxin-gene catalogue generated from a novel *V. berus* genome assembly. In parallel, we assayed general protease and PLA_2_ activities, as well as FXa-, thrombin-, and plasmin-like activities, and cytotoxicity toward mammalian cell lines. We identified two distinct age-related venom phenotypes (ontotypes): an svMP/CTL-rich ontotype A (≤1 year) and an svSP/PLA_^2^_-rich ontotype B (≥2 years). Functionally, protease activity decreased with age, whereas thrombin-like, plasmin-like and PLA_2_ activities, and cytotoxicity, increased. Our findings indicate an ontogenetic shift in composition and activities of *V. berus* venom that parallels dietary transitions and potentially reflect adaptation to differing prey physiologies.

## 2 Introduction

Venom is an ecologically relevant trait that has evolved independently more than 100 times across the animal kingdom ^1–3^. Variation in venom composition and bioactivity occurs at all taxonomical levels, driven by various factors but especially by adaptation to differing ecological demands ^2,4^.

In snakes, venoms are known in over 2,500 species and are mainly applied for hunting. Therefore, shifts in prey spectrum and foraging ecology may have a crucial impact on the venom composition ^3–5^. Factors like origin locality of individuals, sex, and an ontogenetic shift in prey preferences play a role in determining intraspecific venom variation ^6–9^. Along a dietary shift during ontogeny, changes in venom composition have been extensively described in several Crotalinae ^10–12^, and in some Viperinae with *Daboia russelii* ^13^, *Vipera ammodytes* ^14,15^ and *Vipera latastei* ^16^.

The common adder, *Vipera berus*, has the broadest geographic range of any snake species, ranging from Great Britain to the Pacific coast of Asia, occupying various habitats, different in climatic conditions, habitat structure and prey availability ^17–19^. Similar to other vipers, *V. berus* exhibits a dietary shift during maturation. Juveniles predominantly feed on ectothermic prey, such as amphibians and reptiles, and gradually incorporate endothermic prey such as birds and small mammals, which constitute most of the diet of adult individuals ^20–22^. Due to its widespread distribution, *V. berus* is one of the species most frequently involved in snakebite accidents in Europe. Envenomation by *V. berus* usually causes pain at the bite site, discoloration and oedema formation, as well as nausea, hypotension and diarrhea ^23^. Although severe courses are rare, they may involve coagulopathy, extensive tissue damage, renal failure, seizure and myocardial infarction, and can potentially lead to death. ^24–26^. These severe symptoms usually occur in young, elderly, or otherwise compromised individuals, particularly when medical intervention is delayed or absent.

*Vipera berus* venom is dominated by phospholipases A_2_ (PLA_2_s), snake venom serine proteases (svSPs) and snake venom metalloproteinases (svMPs), as well as C-type lectins including snaclecs/C-type lectin-related proteins (CTLs), L-amino acid oxidases (LAAOs), and cysteine-rich secretory proteins (CRISPs) ^27–31^. Early findings by Nedospasov and Rodina (1992) report a marked age-related shift in serine protease (thrombin- and kallikrein-like) activity in *V. berus* venom, increasing sharply from the first year of life towards older age groups. Furthermore, Malina *et al*. (2017) identified higher molecular weight components by SDS-PAGE in Hungarian juvenile *V. berus* specimens compared to the adults. However, comprehensive insights into the extent of intraspecific venom variation along maturation is pending.

In the present study, we investigated *V. berus* venom from early life (pre-first year) through to the adult stage (≥4 years of age). Individual venoms were pooled according to the age groups and profiled using SDS-PAGE and genome-guided shotgun proteomics. Furthermore, we assessed enzyme activity for the PLA_2_ and proteases, evaluated Factor Xa (FXa)-like, thrombin-like and plasmin-like activity, and tested cytotoxicity on two different mammalian cell lines (i.e., MDCK II and Calu-3). Our work provides a compositional and functional perspective on the intrinsic dynamics of the venom of Earth’s most widespread snake.

## 3 Materials and Methods

### 3.1 Venom

Venoms of 30 captive-bred *V. berus* individuals, originating from northern Bavaria (Germany) were donated by members of the “Terrarienclub Bayreuth und Umgebung e.V.” and stored on dry ice upon lyophilisation. Lyophilised venoms were weighed and pooled according to the age of the specimens, which were classified as Neonate (newborn), Yearling (1 year), Juvenile (2 years), Subadult (3 years), and Adult (≥4 years). For each age group, venoms of six individuals were redissolved in double-distilled water (ddH_^2^_O) and combined in equal dry-mass proportions. Lyophilised aliquots were stored upon utilisation at –20 °C. Each specimen’s body size, body-weight and dry venom amount are reported in Supplementary Table S1.

### 3.2 Compositional venom profiling

#### 3.2.1 Reducing and non-reducing SDS-PAGE

Reduced and non-reduced sodium dodecyl-sulfate polyacrylamide gel electrophoresis (SDS-PAGE) with 5 µg of each sample was performed as previously described ^34^. The raw gel image is provided in Supplementary Figure S1. Band densitometry was assessed using ImageJ v1.54d ^35^ and values are provided in Supplementary table S2.

#### 3.2.2 Shotgun proteomics

The proteomic assessment was adapted from our previously established workflow for Shotgun Liquid Chromatography–Electrospray Ionization–Mass Spectrometry (LC-ESI-MS) of animal venoms ^36,37^. For complete denaturation, lyophilised venoms were dissolved to a final concentration of 1.7 µg/µl in an aqueous solution of 6 M Guanidinium hydrochloride (GdnHCl) and 100 mM Tris(hydroxymethyl)aminomethane hydrochloride (Tris/HCl, pH 8.5). Reducing disulfide bonds, venoms were transferred to Protein LoBind tubes (0030108116, Eppendorf) with 10 mM Dithiothreitol (DTT) and were incubated for 30 min at 37 °C, followed by the alkylation of free thiols with 40 mM chloroacetamide, incubated for 30 min at 22 °C in the dark. Initiating protein digestion, Trypsin/LysC mix (V5071, Promega) was added at a 1:50 ratio (protease-to-protein) and incubated for 1 hour at 37°C and 500 rounds per minute (RPM) shaking. In order to reactivate trypsin and to decrease the GdnHCl concentration below 1 M, samples were further diluted 1:7 with 50 mM Tris/HCl (pH 8.0), continuing the incubation overnight at 37 °C and 500 RPM shaking. The next day, the digestion was stopped by adding 1.5 % trifluoroacetic acid and the resulting peptides were cleaned-up and desalted using C18-Chromabond columns (730011, Macherey-Nagel). The eluted peptides were dried in a vacuum-concentrator plus (Eppendorf) and again redissolved in 20 µl of an aqueous solution with 5% (v/v) MeCN and 0.15% (v/v) HFo by vortexing. Transferred to a 96-well PCR plates (PCR-96-FS-C, Axygen), the peptides were sonicated for 5 min in a water bath, and loaded for LC-ESI-MS/MS analysis.

Upstream LC of 5 µl redissolved peptides was performed on an UltiMate 3000 RSLCnano device (Thermo Fisher Scientific) with a PepMap Neo Trap column (Thermo Fisher Scientific) for concentration and desalting, and a 50 cm µPAC column (PharmaFluidics, Thermo Fisher Scientific) for separation. The analytical column was kept in an oven at 35°C throughout the analysis and the peptide elution was performed using a linear gradient of buffer A (HPLC-grade H_2_O with 0.1 % (v/v) HFo) and buffer B (MeCN with 0.1% (v/v) HFo) at flow rate of 0.7 µl/min, given at min (%B): 0-90 (5-35%), 90-100 (35-85%), 100-108 (cons. 85%), and 108-118 min (cons. 5%) for re-equilibration. The peptide elution was directed to a TriVersa NanoMate (Advion) robot for chip-based nano electrospray ionization, applying a 1.7 kV electrospray with a source temperature of 250 °C. Generally, MS analysis of the peptides was carried out on an Orbitrap Eclipse Tribrid MS (Thermo Fisher Scientific) in positive mode and MS2 spectra were obtained in data-dependent acquisition (DDA) with CID fragmentation. Data acquisition and analysis was done with Xcalibur v4.3.73.11. (Thermo Fisher Scientific).

Using novel genome data of *V. berus* (NCBI: BioProject PRJNA1391388, BioSample SAMN54603525) including two haplotypes, we generated a toxin gene catalogue (see Supplementary Table S3) applying ToxCodAn-Genome ^38^, a dedicated bioinformatic pipeline designed for the annotation of toxin-coding regions in the genomes of venomous taxa. The tool was run in a Conda v25.9.1 ^39^ environment and initialized according to the authors’ instructions (set-up details in Supplementary Table S3). Toxin gene identification was performed in default options on both haplotypes with the tool’s internal databases “Viperidae” and “Elapidae”. Subsequently, signal peptides were removed from the final annotation files via SignalP v6.0 (Organism: Eukarya; Model Mode: Slow) ^40^, and duplicate sequences deleted using seqkit v2.10.1 ^41^ in default options. The protein sequences were annotated via InterProScan v5.66-98.0 ^42^ as well as ultra-sensitive DIAMOND v2.1.13 ^43^ searches against two databases: UniProtKB/Swiss-Prot 2025_02 ^44^ and VenomZone ^45^. In each search, we retrieved the best hit for highest sequence similarity, query coverage and lowest e-value, respectively. The E-value was set to a maximum of 1×10^−3^ and max-target-seqs was set to 0 to search the entire database. Similarity was calculated with the BLOSUM62 matrix ^46^ using BioPython v1.85 ^47^. The resulting top DIAMOND hit was used for further analysis. Based on all information collected, sequences were assigned to toxin families.

Protein identification was performed with PEAKS 12.0 (build 20250725, Bioinformatics Solution Inc.) against the generated *V. berus* toxin gene catalogue. Briefly, the following parameters for protein annotation were applied: 15 ppm precursor ion mass tolerance; 3 missed cleavages; carbamidomethylation as fixed modification; 0.5 Da fragment ion mass tolerance in linear ion trap MS2 detection; 0.1 false discovery rate. For the qualitative analysis, we only considered identified protein groups with at least a –10lgP score of 15 and two unique peptides. Annotated proteins were further clustered into protein groups by PEAKS 12.0, based on high sequence similarity, represented by a single “TOP” protein for each protein group. Protein groups were manually curated to account for sample-dependent differences in unique peptide coverage that could otherwise bias NSAF-based quantification. To estimate the relative abundance of the identified toxins, spectral abundance factors (SAFs) were calculated for each protein group (k) using spectral counts (SpC) for the corresponding “TOP” protein as a proxy for abundance and its protein length (L). We then normalized each SAF (NSAF) by dividing it by the sum (N) of all SAFs within the investigated age group ^48,49^:

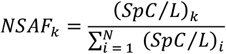

Detailed settings for MS analysis and protein annotation are provided in Supplementary Table S4. The full list of annotated proteins is available in Supplementary Table S5. All RAW MS files have been deposited in the Zenodo repository under the project “DATASET - Mass Spectrometry - Snake venom proteomics of *Vipera berus* (Age Variation); 10.5281/zenodo.17815205”.

### 3.3 Bioactivity profiling

Bioactivity profiling of the samples was based on previously established protocols for viperine snake venoms and applied assessing enzyme activity for proteases, PLA_2_, FXa-like, thrombin-like, and plasmin-like activities ^37,50,51^, as well as effects on mammalian cell viability ^52–54^. All analyses were performed in at least triplicate. For signal detection a Synergy H4 Hybrid Microplate Reader (BioTek) operated with the Gen 5 v2.09 software (BioTek) was used. Unless otherwise stated, the respective measured venom signals (SA) at specific wavelengths were averaged, background-corrected by subtracting the negative control (NC), and normalized against the positive control (PC) as follows, with relative activity (RA, in %) and the mean absorbance/emission 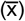:

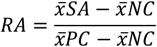

Protease- and PLA_2_ activity assays were conducted in two sperate batches. In addition to the respective positive and negative control, both batches contained the dilution series of the Adults venom sample, which served as a batch reference standard. To enable comparison of results between batch b and batch b, both, the relative activity and the corresponding standard deviation of each diluted sample in batch b, was corrected (c) for technical variance. For each dilution, the relative activity (RA) and standard deviation (SD) of the diluted samples in batch b were multiplied by a correction factor derived from the ratio of the Adults sample’s RA in batch b and batch a:

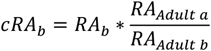

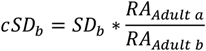

To estimate the standard deviation of the adult sample across batches, an adaptation of the pooled standard deviation formula for independent groups ^55^ was applied with number of replicates in respective batches (n):

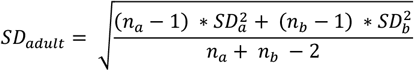

#### 3.3.1 Protease activity assay

The non-specific Protease Activity Assay Kit (Calbiochem, cat. no. 539125) was used, as previously described ^34^. Venoms were redissolved (Reaction Buffer 6:1 ddH_2_O) and added at a final concentration of 100 μg/ml, similar to the negative control (Reaction Buffer 6:1 ddH_2_O, 0%) and positive control (166 µg/ml trypsin in Reaction Buffer 6:1 ddH_2_O, 100%). The samples were incubated for 2 h at 37 °C, shaking at 120 rpm on a Multitron device (Infors HT) following the absorption measurement at λ = 492 nm. Raw data from the protease activity assay is provided in Supplementary Table S6.

#### 3.3.2 Factor Xa-like activity assay

The Factor Xa (FXa)-like activity was investigated using the FXa Activity Fluorometric Assay Kit (MAK238-1KT, Sigma-Aldrich), as previously described ^51^. Venoms were redissolved (Factor Xa Assay Buffer 20:1 ddH_2_O) and added to the substrate at a final concentration of 12.5 μg/ml, similar to the negative control (FXa Assay Buffer 20:1 ddH_2_O, 0%) and the positive controls (2 µg/ml FXa Enzyme Standard in FXa Assay Buffer 20:1 ddH_2_O, 100%). The assay was incubated for 10 min at 37 °C, protected from light, following signal measurement with excitation set to λ = 350 nm and fluorescence detection at λ = 450 nm. Raw data from the FXa-like activity assay is provided in Supplementary Table S7.

#### 3.3.3 Thrombin-like activity assay

The thrombin-like activity was investigated using the Thrombin Activity Fluorometric Assay Kit (Sigma-Aldrich, MAK242, as previously described ^51^. Venoms were redissolved (Thrombin Assay Buffer 20:1 ddH_2_O) and added to the substrate at a final concentration of 12.5 μg/ml, similar to the negative control (Thrombin Assay Buffer 20:1 ddH_2_O, 0%) and the positive control (0.3 µg/ml Thrombin Enzyme Standard in Thrombin Assay Buffer 20:1 ddH_2_O, 100%). The assay was incubated for 10 min at 37 °C, protected from light, following signal measurement with excitation set to λ = 350 nm and fluorescence detection at λ = 450 nm. Raw data from the thrombin-like activity assay is provided in Supplementary Table S8.

#### 3.3.4 Plasmin-like activity assay

The plasmin-like activity was assessed using the Plasmin Activity Assay Kit (MAK244, Sigma-Aldrich, as previously described ^51^. The lyophilised venoms were redissolved (Plasmin Assay Buffer 20:1 ddH_2_O) and added to the substrate at a final concentration of 12.5 μg/ml, similar to the negative control (Plasmin Assay Buffer 20:1 ddH_2_O, 0%) and the positive controls (5 µg/ml Plasmin Enzyme Standard in Plasmin Assay Buffer 20:1 ddH_2_O, 100%). The assay was incubated for 10 min at 37 °C, protected from light, following the absorption measurement at λ = 405 nm. Raw data from the plasmin-like activity assay is provided in Supplementary Table S9.

#### 3.3.5 Phospholipase A_2_ activity assay

Phospholipase A_2_ activity was assessed using the EnzChek Phospholipase A_2_ Assay Kit (Invitrogen, cat. no. E10217), as previously described ^51^. Venoms were redissolved (Reaction Buffer 20:1 ddH_2_O) and added at a final concentration of 12.5 μg/ml to the substrate, similar to the negative control (Reaction Buffer 20:1 ddH_2_O, 0%) and positive control (5 U/ml purified bee venom PLA_2_ in Reaction Buffer 20:1 ddH_2_O, 100%). After incubation for 50 min at room temperature, signals were detected at λ = 515 nm following excitation at λ = 470 nm. Raw data from the phospholipase A_2_ activity assays is provided in Supplementary Table S10.

#### 3.3.6 Cell viability assay

The effect of venoms on the cell viability of Madin-Darby canine kidney II (MDCK II; kindly provided by Prof. Dr. Eva Böttcher-Friebertshäuser, Institute of Virology, Philipps University, Marburg) and human epithelial lung adenocarcinoma (Calu-3; SCC438, Merck) cell lines was assessed using the CellTiter-Glo Luminescent Cell Viability Assay (G7570, Promega), as previously described ^52^. The venoms were redissolved in cultivation medium and added to confluent cells at a final concentration of 12.5 µg/ml, similar to the negative control (ddH_2_O, 100% cell viability) and the positive control (100 µM ionomycin in DMSO, 0% cell viability). Following incubation for 48 h at 37 °C in a 5% CO2 atmosphere, luminescence was measured according to the manufacturer’s protocol. Raw data from the cell viability assay is provided in Supplementary Table S11.

## 4 Results

### 4.1 Morphometrics across ontogeny

All 30 individuals *V. berus* were measured for body size and mass, milked and the dried venom yield weighed after lyophilisation. Mean values (Table 1) were comparable for Neonates and Yearlings, and increased over Juveniles to Subadults and Adults. The Subadults were larger and heavier than Juveniles, but had comparable dried venom yields. Furthermore, the Subadults and Adults had a comparable mean body size, whereas Adults showed a higher mean body mass and a greater dry venom yield compared to Subadults.

**Table 1.** Metrics: The investigated individuals of each age group (Neonate (0 years), Yearling (1 year), Juvenile (2 years), Subadult (3 years), Adult (≥4 years)) were measured for their body size (cm), body mass (g) and the amount of dry venom provided (mg). Displayed are mean values ± standard deviation (n = 6).

| Metric | Neonate | Yearling | Juvenile | Subadult | Adult |
| --- | --- | --- | --- | --- | --- |
| Body size (cm) | 22.0 ± 1.7 | 21.7 ± 3.2 | 34.7 ± 4.2 | 53.1 ± 5.5 | 56.3 ± 3.1 |
| Body mass (g) | 9.2 ± 2.7 | 7.7 ± 2.3 | 23.0 ± 7.6 | 99.7 ± 42.3 | 120.8 ± 59.0 |
| Dried venom (mg) | 0.73 ± 0.26 | 0.87 ± 0.22 | 1.88 ± 0.71 | 2.05 ± 0.28 | 4.75 ± 3.21 |

### 4.2 Compositional venom profiling

After pooling, 5 µg of venom from each of the five age groups was analysed by reducing and non-reducing SDS-PAGE (Figure 1 A & B). Under reducing conditions (Figure 1 A), band patterns were observed from 12-70 kDa across all samples. Overall, Neonates and Yearlings exhibited highly similar venom profiles, as did Juveniles, Subadults and Adults. The main differences between these groups occurred at 35-37, 31-33, and 14-15 kDa, which were weaker in younger age groups; at 50-60 kDa, a double band at the younger age groups that transitions to a more intense band at 60 kDa band in older stages; and at 65-70 kDa which was strong in Neonates and Yearlings but nearly absent from Juveniles onwards.

**Figure 1:**
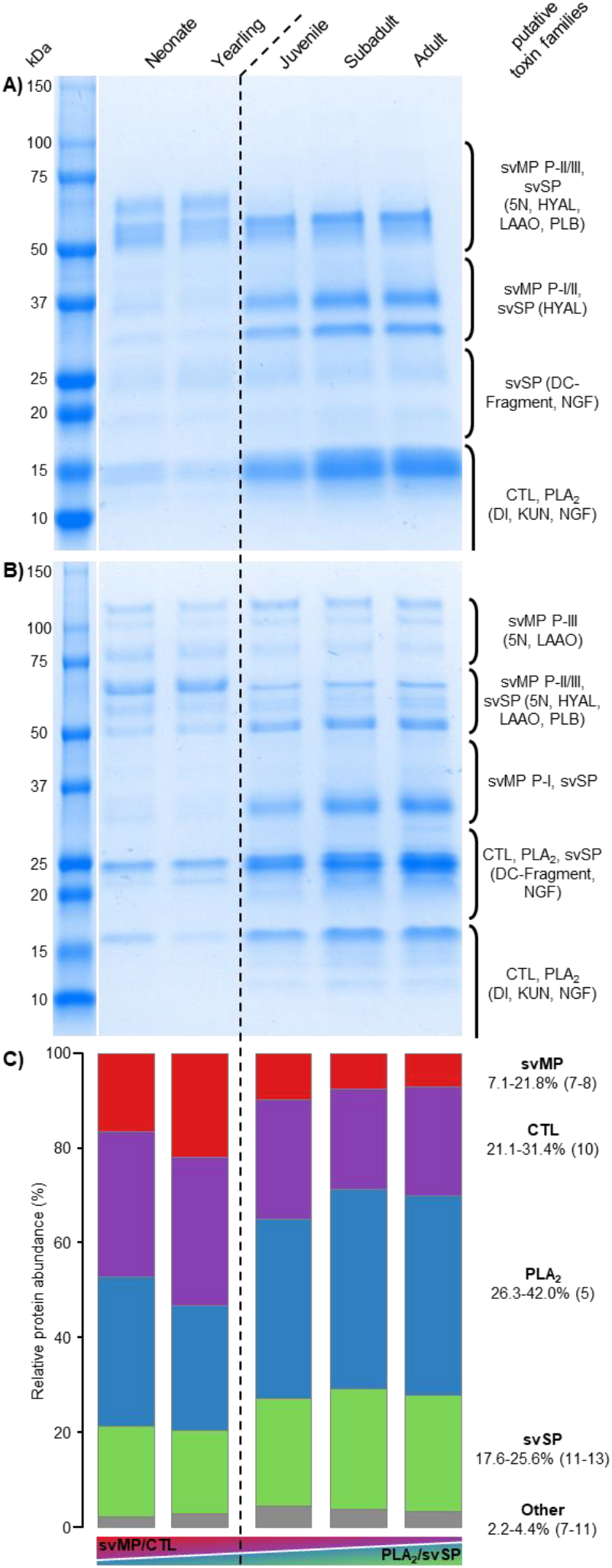
Venom composition profiling across age groups in *Vipera berus*. Pooled single venoms (n = 6) representing five ontogenetic stages (Neonate (0 years), Yearling (1 year), Juvenile (2 years), Subadult (3 years), Adult (≥4 years)) were analyzed by A) reducing SDS-PAGE (5 µg), B) non-reducing SDS-Page (5 µg), and C) shotgun proteomics. Putative toxin families were assigned based on the molecular masses reported in previous *V. berus* venom proteomic studies ^27–29,31^. Major toxin families (sensu ^56^): snake venom serine protease (svSP), snake venom metalloproteinase (svMP; subfamilies P-I, P-II/disintegrin (DI), P-III/DI-like-cysteine-rich (DC)-Fragment), phospholipase A_2_ (PLA_2_), C-type lectin including snaclec/C-type lectin-related protein (CTL). Other toxin families: 5′-nucleotidase (5N), type-B carboxylesterase/acetylcholinesterase (ACHE), chitinase, cystatin (CYS), ficolin, hyaluronidase (HYAL), Kunitz-type serine protease inhibitor (KUN) L-amino acid oxidase (LAAO), lipase (LIP), nerve-growth factor (NGF), phospholipase B-like (PLB), phosphodiesterase (PDE). Number in brackets indicate the count of identified proteins and the dashed line indicates a turnover in venom composition between the age groups.

Under non-reducing conditions (Figure 1 B), band patterns ranged 12-120 kDa. As seen under reducing conditions, the two youngest and three oldest presented highly similar venom profiles, respectively. In Neonates and Yearlings, bands at 75-80 and 65-70 kDa were more intense, whereas older age groups showed stronger bands at 115-120, 50-52, 31-36, 24-26, and 16-17 kDa. The 55-60 kDa band transition from broad band in the two youngest to a distinct double band in older age groups. Differences between Neonates and Juveniles were minor (Neonates > Yearlings: 115-120, 100-105, 31-36 kDa; Neonates < Yearlings: 20-22 kDa). A 29-30 kDa band was unique for Adults, while the bands at 12-15 kDa appeared exclusively from Juveniles onwards. The region at around 20-23 kDa showed a more complex pattern across age groups: a faint single band in Neonates, more intense in Yearlings, then from Juveniles onwards transitioning into an increasingly more intense double band, with the upper portion merging into the 24-26 kDa band and the lower fading near 20 kDa.

Shotgun proteomics was performed on pooled venoms representing the five age groups (Figure 1C). Equal concentrations of pooled venoms (1.7 µg/ml) were used for all samples and toxin-specific annotation was applied uniformly Peptide spectra were annotated, assigned to protein groups by PEAKS 12.0 based on and quantified using a toxin specific database derived from a genome of *V. berus*, comprising 81 toxin gene candidates from two *V. berus* haplotypes. In total, 52 toxin protein groups were identified across all age groups, ranging 40-45 protein groups detected per age group and 39 protein groups shared among all.

The normalized SAF (NSAF) per age group showed that the major venom components svMP, CTL, PLA_2_ and svSP (sensu ^56^) together comprised 95.6-97.8% of relative toxin abundance. The svMP and CTL, as often forming svMP P-III multimers, show their highest combined proportions in the venom of Neonates (47.1%: 16.5% svMP, 30.6% CTL) and Yearlings (53.2%: 21.8% svMP, 31.4% CTL), and decrease towards Juveniles (34.9%: 9.6% svMP, 25.2% CTL), Subadults (28.7%: 7.5% svMP, 21.1% CTL) and Adults (30.1%: 7.1% svMP, 23.0% CTL). In Contrast, the combined proportions of PLA_2_ and svSP increased steadily from Neonates (50.7%: 31.5% PLA_2_, 19.2% svSP) and Yearlings (43.9%: 25.5% PLA_2_, 19.3% svSP), towards Juveniles (60.7%: 37.8% PLA_2_, 22.9% svSP), Subadults (67.5%: 41.9% PLA_2_, 25.6% svSP), and Adults (66.6%: 42.0% PLA_2_, 24.6% svSP).

The other toxin families contributed 2.2-4.4% per age group. Kunitz-type inhibitors (KUNs), nerve growth factors (NGFs), phospholipases B-like (PLBs), cystatins (CYSs), LAAOS, and 5’-nucleotidases (5Ns) were present in all age groups. Hyaluronidases (HYALs), lipases (LIPs), type-B carboxylesterase/acetylcholinesterases (ACHEs), ficolins, and chitinases occurred intermittently.

Taken together, both compositional analyses highlight a consistent age-related pattern in V. berus venom compositions. Neonates and Yearlings display highly comparable profiles, as did Juveniles, Subadults, and Adults, with pronounced compositional differences occurring between the first and second year of life. Early-life venom profiles are characterized by higher-molecular weight components and a higher relative abundance of svMP and CTL, whereas later-life venom profiles are characterized by an increase in lower-molecular weight components and were dominated by the relative abundance of PLA_^2^_ and svSPs.

### 4.3 Bioactivity profiling

After characterizing the proteome compositions, the five age groups were assayed for protease activity (100 µg/ml) and PLA_2_ activity, FXa-, thrombin- and plasmin-like activities, as well as effects on the viability of MDCK II and Calu-3 cells at 12.5 µg/ml (Figure 2).

**Figure 2:**
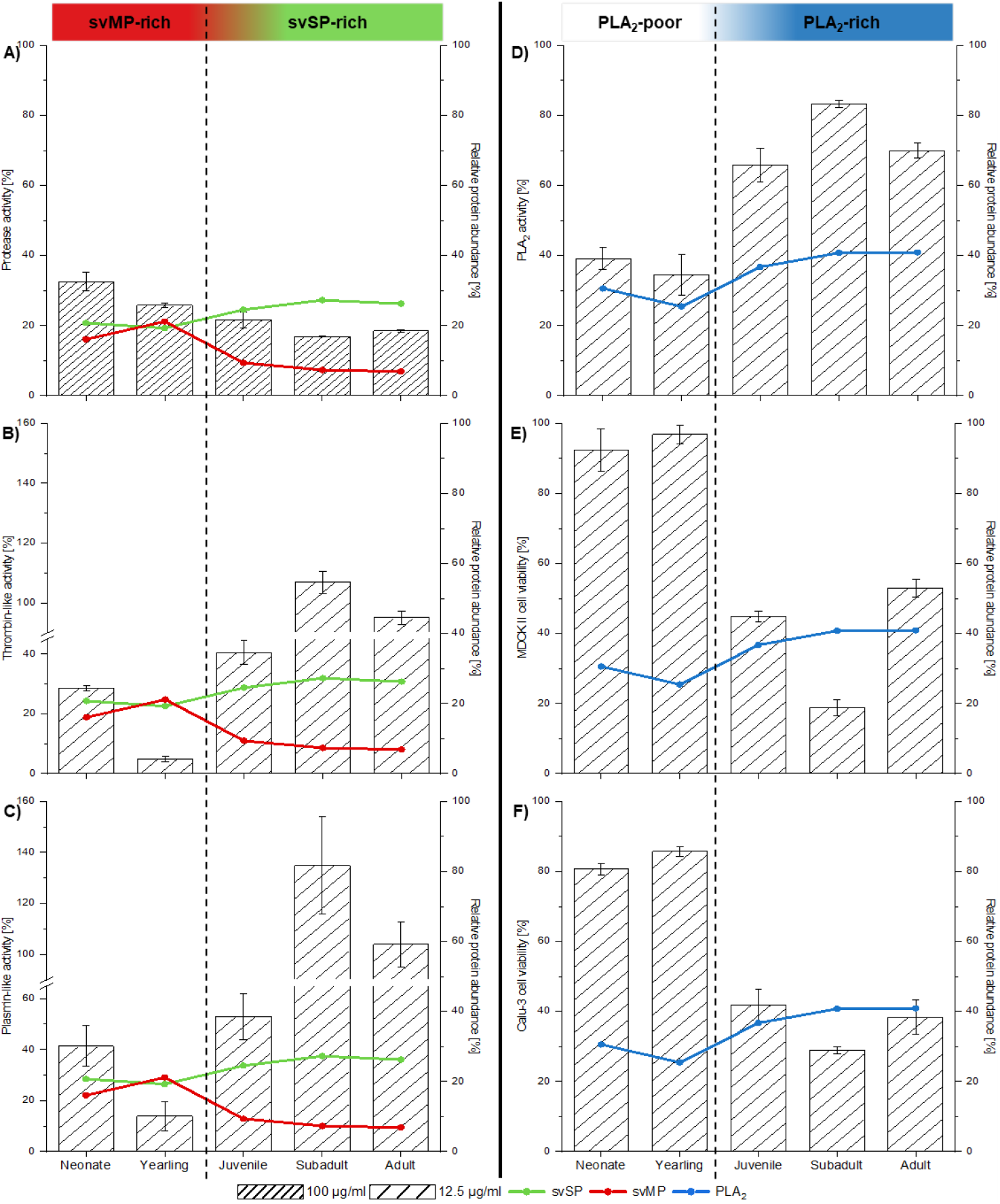
Venom bioactivity profiling along relevant protein diversity across age groups in *Vipera berus*. Age-related venom pools (n = 6 per age groups Neonate (0 years), Yearling (1 year), Juvenile (2 years), Subadult (3 years), Adult (≥4 years)) were applied at 100 µg/ml for A) protease activity, and at 12.5 µg/ml for B) thrombin-like activity, C) plasmin-like activity, D) phospholipase A_2_ (PLA_2_) activity, and cell viability of E) MDCK II and F) Calu-3 cells. Values represent the averaged activity of technical replicates (n = 3), base-line corrected against the respective negative control and normalized to the respective positive control (± SD). The secondary Y-axis presents the relative abundance of relevant toxin families across the age groups. The dashed line indicates a turn-over in general bioactivity between the age groups.

Relative protease activity decreased with age from 33% to 17% (Figure 2A). Thrombin-like activity was detected across all age groups and ranged from 5% to over 100% (Figure 2B). In contrast to the general protease activity, thrombin-like activity decreased from Neonates to Yearling before increasing with age, peaking in Subadults. Plasmin-like activity presented a similar activity pattern, ranging from 14% to over 100% (Figure 2C). Factor Xa-like activity was absent in all age groups (see Supplementary table S7).

Relative PLA_2_ activity ranged from 35% to 83 % and showed a pattern comparable thrombin-like and plasmin-like activity, although Juveniles exhibited activity levels more similar to the older age groups than to the younger ones (Figure 2D). Treatment of MDCK II cells (Figure 2E) and Calu-3 cells (Figure 2F) with the age-related pooled venoms reduced cell viability in a comparable manner (MDCK II: 19-97%; Calu-3: 29-86%). Venoms from Neonates and Yearlings had little effect, whereas older age groups strongly reduced cell viability, strongest effect observed with Subadults’ venom.

Similar to the pattern observed in the compositional analysis, the venom of Neonates and Yearlings present comparable bioactivities, as do Juveniles, Subadults, and Adults. Overall, the early-life venoms (Neonates and Yearlings) presented higher general protease activity compared to the later-life venoms (Juveniles, Subadults, and Adults), while the latter exhibit higher PLA_2_, Thrombin-like and Plasmin-like activities and have a strong impact on the cell viability at the same concentrations.

## 5 Discussion

Identifying the drivers of snake venom diversity at both inter- and intra-specific levels is essential for understanding snake venom evolution and for improving snakebite treatment ^9^. Intraspecific venom variation has frequently been associated with dietary changes due to differences in locality, environment, or the snake’s morphology, associated for instance with sexual dimorphism and ontogeny ^6,8,12^. In snakes, the size of ingestible prey is limited by a snake’s mouth-gape, which is positively correlated with the snake’s overall body size ^57,58^. Consequently, as younger individuals typically feed on smaller prey items than older, larger individuals, venom variation between different age classes is expected as an adaptation to favour the effective subjugation of different prey.

In our study, both compositional analyses as well as bioactivity assessments reveal two comparable age-related venom ontotypes. Neonates (newborn) and Yearlings (one-year-old) hereafter referred to as ontotype A, share similar venom profiles, as do Juveniles (two-year-old), Subadults (three-year-old), and Adults (≥ four-year-old), collectively referred to as ontotype B. The distinction between these ontotypes is manifested by a shift from an early-life venom ontotype A dominated by higher-molecular weight components (>50 kDa; SDS-PAGE) and exhibiting approximately equal relative abundance of svMP and CTL to PLA_2_ and svSP (genome-guided shotgun proteomics) to a later-life venom ontotype B. This ontotype B shows an increased representation of lower-molecular weight components (>50 kDa; SDS-PAGE) and the relative abundance of major venom components shifted towards PLA_2_ and svSP (genome-guided shotgun proteomics). While SDS-PAGE profiling indicates a sharp shift in venom composition between one- and two-year-old individuals, the observed changes in relative toxin abundance (genome-guided shotgun proteomics) and functional bioactivity appears more gradual from the first to the third year of living.

In line with the findings of Malina *et al*. (2017), venom ontotype A largely comprises higher-molecular-weight components (>50 kDa), whereas venom ontotype B shows a notable increase in 30-37 kDa and lower-molecular-weight (13-26 kDa) bands. Based on previous two-step quantification approaches performed on *V. berus* venom ^27–30^ bands >50 kDa correspond to svMPs (P-III) and LAAOs. The 20-50 kDa bands generally comprise svMPs, svSPs, and PLA_2_s. The conspicuous 31-37 kDa bands are usually comprised by svSPs, and the 20-25 kDa bands typically represent dimeric PLA_2_s. Upon reduction, the monomeric PLA_2_ migrate into a strong 13-15 kDa band. Lower bands (<20 kDa) usually contain CTLs, PLA_2_s, and DIs. Our genome-guided shotgun proteomics presented svMP, CTL, PLA_2_, and svSP as high abundant venom components, largely consistent with previous studies ^27–31^. Compared with these studies, our results show less diversity among toxin families, and varying proportions of identified toxin families in *V. berus* venom can already be observed in those studies (e.g. svMP range from 0.2 to 19%, PLA_2_ from 10 to 59%, LAAO from 0 to 34%).

Within the genus *Vipera*, an ontogenetic dietary shift from an ectotherm-to an endotherm-based diet has been described along a transition from a svMP-rich venom in younger individuals to a svSP/PLA_2_-rich venom in older ones ^14–16^, although the timing of that transition remained unclear. In *V. berus*, early work by Nedospasov and Rodina (1992) reported increasing thrombin- and kallikrein-like activities from one year of age onwards, suggesting corresponding changes in venom composition. Concordantly, our quantitative genome-guided shotgun proteomics and bioactivity profiling corroborated the SDS-PAGE results, highlighting a shift in *V. berus* venom from a ratio of approximately 1:1 to 2:1 of PLA_2_ and svSP to svMP and CTL from the age of two years old onwards (Figure 2). In parallel, general protease activity steadily decreased from Neonates onwards, which is in line with the general decrease in relative svMP as well as the overall protease (svMP and svSP) abundance. An exception is Yearlings’ venom, with its protease activity decreasing, compared to Neonates, while protease abundance increases. However, this might be due a putatively lower absolute abundance of toxins in the investigated sample as indicated by SAF-value of Yearlings venom being the lowest of all age groups (see Supplementary table S5). Because the assay employed FTC-Casein, a substrate with multiple cleavage sites, the activity of the non-specific svMP ^59^ is likely to be disproportional displayed compared to the more selective svSP ^60^. The similar venom compositions and bioactivities in Neonates and Yearlings might be correlated to similar morphological and physiological constraints of ingestible prey ^57,58^. Ectothermic prey available to young *V. berus* is typically smaller in size than endothermic prey, and possess a slower metabolism and lower heart rates ^61^, factors that likely reduce systemic venom distribution. Ontotype A venoms, which are rich in svMPs, promote severe tissue-degradation and systemic hemorrhage, ^59,62^, and may therefore improve hunting success on ectotherms for younger *V. berus*, possibly reducing the chances of retaliation by the prey, and/or aiding digestion.

In the venom of Juveniles, relative svSP abundance increases while the relative svMP abundance shows a marked decrease, marking the pivotal stage between venom ontotype A and venom ontotype B. This is further supported by venom profiles and the activity assays. Correspondingly, the thrombin- and plasmin-like activities of ontotype B increase compared to venom ontotype A. Although these assays are designed to investigate thrombin/plasmin-like activities, the chromogenic substrates in these assays, mimicking the cleavage site of physiological thrombin and plasmin, can also be cleaved by more broadly specific proteases, potentially inflating the observed coagulation-specific activity. In addition to the slight increase of svSP-driven activity, PLA_2_ activity drastically increased in the venom of two years old specimens. Enzymatic PLA_2_s are typical cytotoxic viperid venom components that disrupt cell membranes ^63,64^. Therefore, the observed decrease in viability of the mammalian cell lines MDCK II and Calu-3 align with the increase in PLA_2_ activity.

The age-related change in *V. berus* venom composition and bioactivity coincides with an increase in mean body size of approximately 60% in Juveniles compared to younger specimens (see Table 1). This putatively overcomes the initial gape size limitations and enables the predation of larger prey, such as small endotherms, offering a higher energetic gain ^57,58^. While being in a transition state in Juveniles, in the older Subadults and Adults, venom ontotype B is fully established, with the compositional profile observed in Juveniles remaining consistent. In addition, the thrombin- and plasmin-like activity strongly increases, whereas PLA_2_ activity and the impact on MDCK II and Calu-3 cell viability increase less. Endothermic prey usually exhibits higher blood pressures and heart rates than ectothermic prey ^61^. In light of this, a venom composition rich in svSP and PLA_2_, which typically disrupt haemostasis and elicit cytotoxicity, might be advantageous for targeting prey with a high metabolic rate, expected to accelerate the spread of toxins in the blood system.

The ongoing debate on diet-driven venom evolution has produced differing hypotheses on how dietary diversity and specialisation shape snake venom phenotypes. Recent studies ^65–67^ generally support the view that snakes with a phylogenetically diverse diet possess more compositionally diverse venoms ^68,69^, reflecting selective pressures to target physiological systems of the varied prey items they feed on. Conversely, species with more specialised diets tend to exhibit less diverse venoms, optimised for efficiency against a narrower range of prey targets ^70^. In this scenario, our findings appear to align with this hypothesis: juvenile *V. berus* display less diverse venoms corresponding to their diet primarily consisting of reptiles and amphibians, whereas adults exhibit greater venom diversity alongside a broader prey spectrum.

Taken together, our data suggest a gradual age-related shift from a younger venom ontotype A to ontotype B in older *V. berus*, with the pivotal stage likely occurring around the second year of development. Our results indicate that an ontogenetic shift involving by two different venom ontotypes in *V. berus* occurs. The age-related transition appears to be gradual resulting in an ‘intermediate’ venom ontotype around the second year of life. A similar ‘intermediate’ venom composition has been observed for *V. latastei* ^16^. In our study, a decrease in general protease activity can be observed in Yearlings, but it still resembles the venom profile of Neonates more than that of venom ontotype B. Moreover, in Juveniles, the venom composition is already resembling venom ontotype B as suggested by SDS-PAGE, but further shifts towards a svSP/PLA_2_ dominated venom with increased cytotoxicity properties in the older Subadults and Adults.

## 6 Conclusion and Outlook

Our data reveal a clear ontogenetic shift in *V. berus* venom composition and bioactivity, from a broad-specific svMP/CTL-rich venom (ontotype A) to a svSP/PLA_2_-rich venom (ontotype B). This is paralleling a dietary transition from smaller, metabolically slow ectothermic prey to larger, metabolically active endothermic prey. The shift in venom composition appears gradually, with a turning point at the second year of life, which coincides with physical growth, suggesting ecological adaptation to different trophic ecologies. While SDS-PAGE profiles and genome-based shotgun proteomics provide a robust insight into compositional changes, a deeper analysis, ideally a quantification of fractionated venom, as with the classical ‘snake venomics’ workflow, would be desirable. Furthermore, bioactivity assessment of protease and PLA_2_ activity assays would be strengthened by the addition of specific inhibitors. Including reptilian and amphibian cell lines in the cell viability assays would provide additional insights into prey-specificity. Finally, the contribution of CTL to long-term cytotoxicity and platelet-aggregation should be explored. Further research into ontogenetic shifts in venom composition in *V. berus*, as well as other medically relevant snakes, is pivotal in particularly for snakebite treatment, the efficacy of existing antivenoms against venoms of younger age groups, and the consideration of such venoms in the development of upcoming antivenoms.

## 7 Acknowledgements

We thank the members of the “Terrarienclub Bayreuth und Umgebung e.V.” for donating the venom samples used in this study. We highly appreciate the intellectual support given by various members of the Venture for Interconnection, Protection, Education and Research in Adders, VIPERA e.V., and the support of Celine Zumkeller and Lilien Uhrig. IA, MD, TL, and LS are funded by the Deutsche Forschungsgemeinschaft (DFG, German Research Foundation; refs. 545040837 (IA), 540833593 (MD), and 505696476 (TL and LS), respectively). JE and KH are supported by the BMFTR (Bundesministerium für Forschung, Technologie, und Raumfahrt) project ASCRIBE (Grant number 01KI2024).

