## Supplementary Figure S1 for "Ontogenetic variation in composition and bioactivity of common adder (*Vipera berus*) venom revealed by genome-guided proteomics and in vitro functional assays"

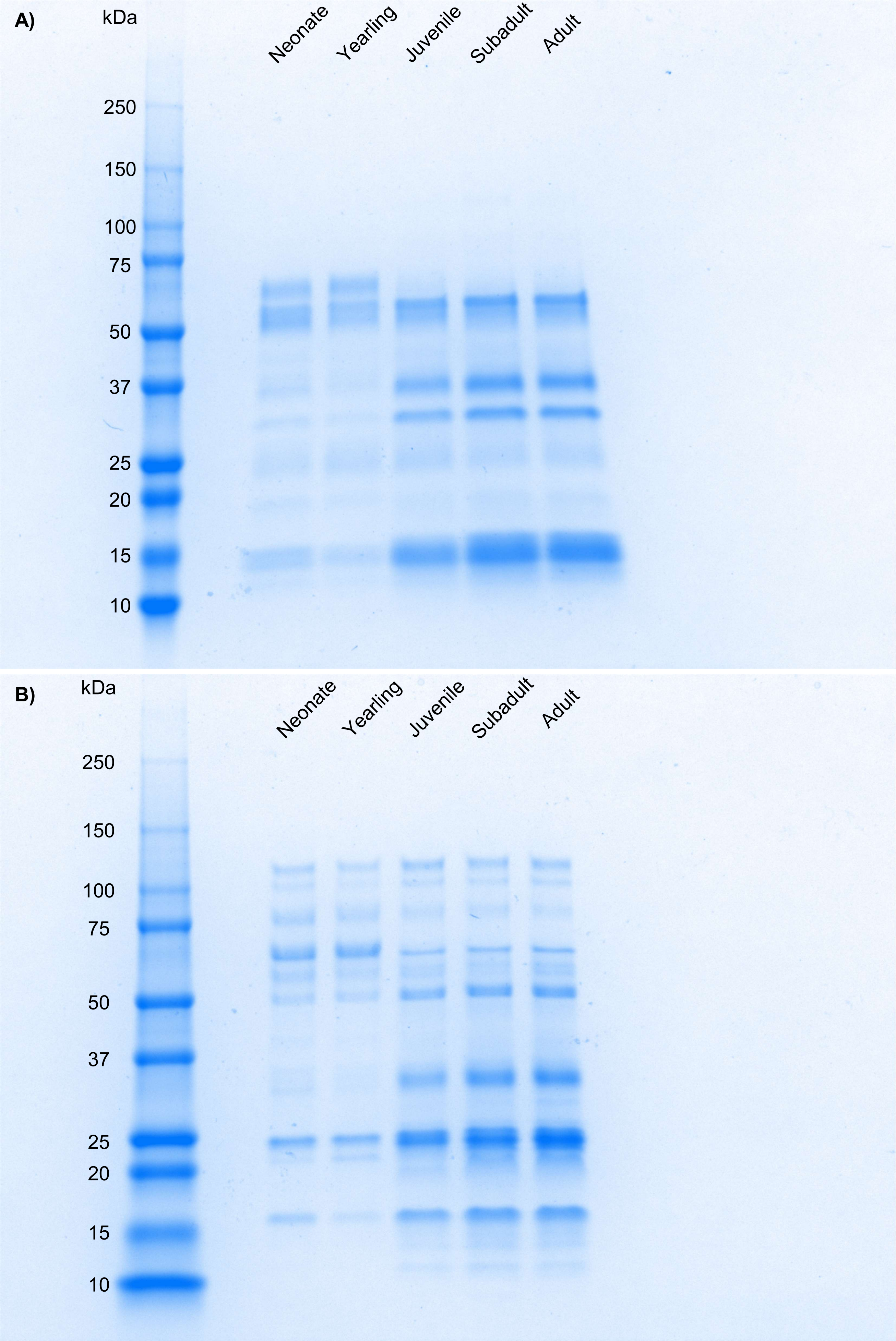


Figure S1: Raw gel images of reducing (A) and non-reducing (B) SDS-PAGE of lyophilised pooled (n = 6) *Vipera berus* venom (5 µg) of five age groups (Neonate: 0 years; Yearling: 1 year; Juvenile: 2 years; Subadult: 3 years; Adult: >4 years).
